## Supplementary figures and images for "Opposing chemosensory functions of closely related gustatory receptors"

### Figure 2-figure supplement 1-Source Data 1

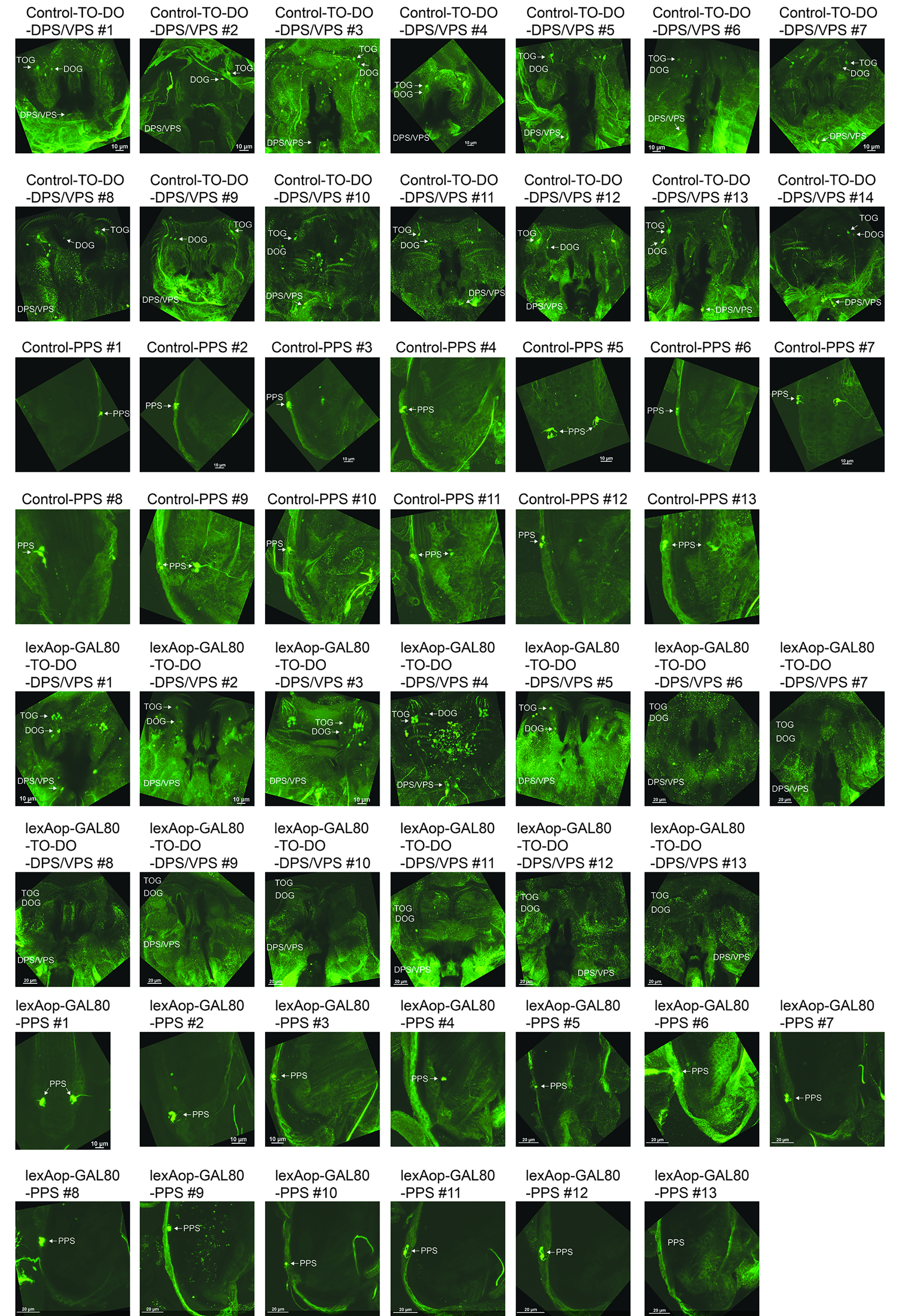
